# Fibroblasts enhance the growth and survival of adult feline small intestinal organoids

**DOI:** 10.1101/2024.03.26.586790

**Authors:** Nicole D Hryckowian, Katelyn Studener, Laura J Knoll

## Abstract

Intestinal organoids are important cell culture models that complement live animal studies of intestinal biology. Adult feline small intestinal organoids are needed for infectious disease research but are difficult to work with due to slow growth and rapid senescence. We introduce a method of co-culturing adult feline small intestinal organoids with growth inhibited human foreskin fibroblast feeder cells to enhance organoid proliferation and survival. With feeder cells, feline jejunal and ileal organoids survived at least nine months in culture until cryopreservation. Fibroblast supplementation also increased the maximum size of cat and mouse intestinal organoids, making them easier for microinjection. The increased proliferation, longevity, and size make fibroblast-supplementation of feline small intestinal organoids a significant improvement on current methods. These methods have high potential to reduce the number of cats used for research and may be applicable for intestinal cells from other animals that are difficult to culture.

## Introduction

The small intestinal epithelium is a single layer of columnar epithelial cells that provide barrier function and absorb nutrients. The epithelium is a crucial component of host defense to orally acquired enteric pathogens like the single-celled eukaryotic parasite *Toxoplasma gondii*. Approximately 30% of humans are chronically infected with *T. gondii* (Rahmanian et al. 2020), and most infections are acquired orally. Oral infection is also key for the *T. gondii* life cycle when it occurs in the definitive feline host. The feline small intestinal epithelium is the only site where parasites undergo sexual reproduction to produce highly infectious and environmentally resistant oocysts. Understanding how *T. gondii* behaves in cat intestine is crucial to interrupting transmission but using live cats in *T. gondii* research is fraught with ethical concerns.

An alternative to studying *T. gondii* in live cats is feline intestinal organoids (enteroids).

Enteroids are derived from intestinal crypt stem cells that divide and differentiate into 3D structures that mimic the cell types and architecture of the intestinal epithelium (Fig. 1). Mouse and human enteroids have enabled important discoveries in infectious disease research, including studies of Apicomplexan parasites (Heo et al. 2018, Wilke et al. 2019, Ramírez-Flores & Knoll 2021). While feline enteroids have been used for infectious disease research by our lab and others, they remain difficult to work with because they undergo early growth arrest and cell death (Powell & Behnke 2017, Tekes et al. 2020).

**Figure 1:**
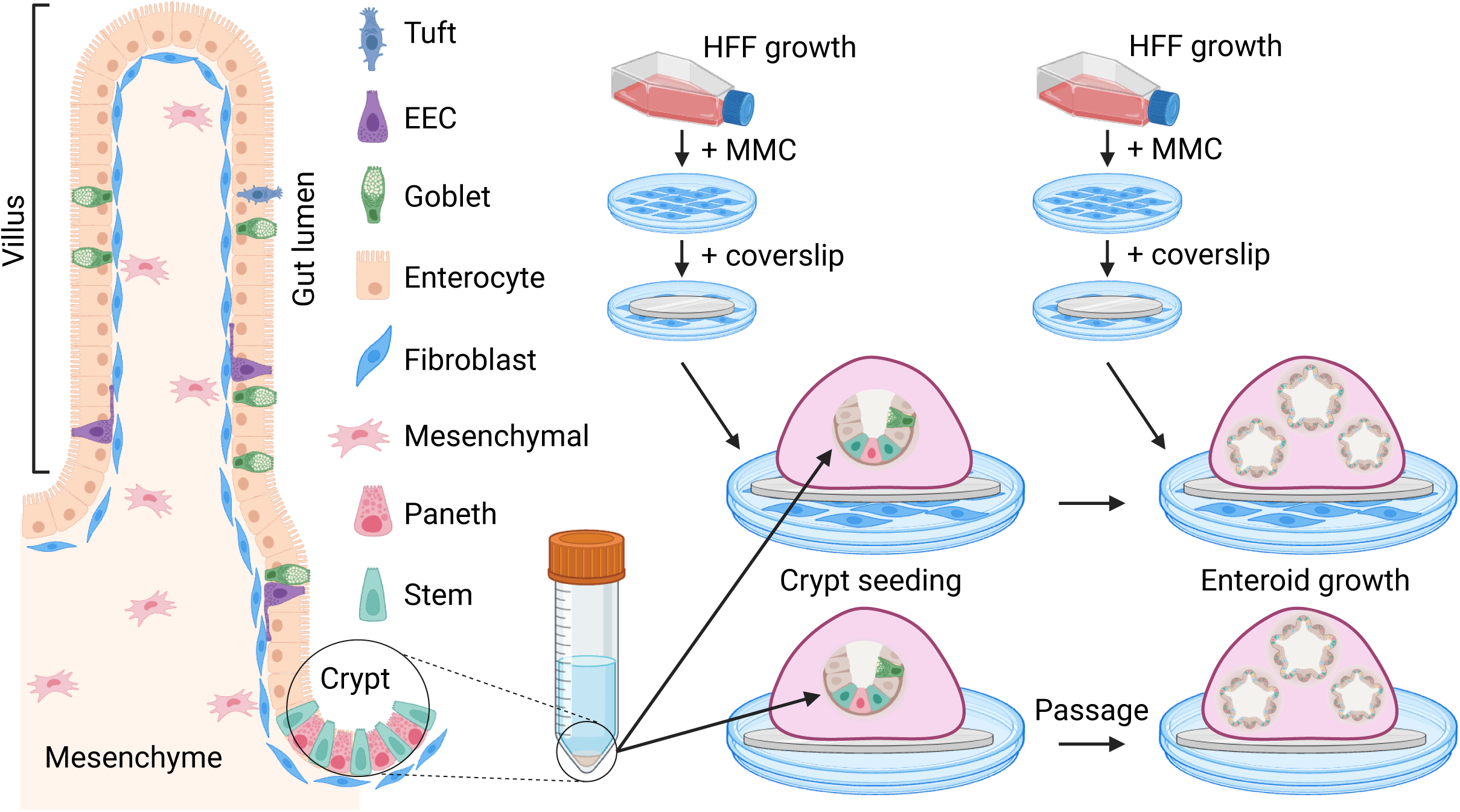
S**m**all **intestinal anatomy and fibroblast supplementation of enteroids.** A schematic of healthy small intestinal epithelium is shown at left, along with the common cell types of the small intestine and gut compartments discussed in this paper. EEC = enteroendocrine. HFF = human foreskin fibroblast. MMC = mitomycin C. A simplified schematic of crypt isolation for enteroid preparation with and without HFFs is shown at right. New HFFs must be prepared prior to each enteroid passage. Image created with Biorender.com.

*In vivo*, intestinal stem cells rely on growth factors from crypt-resident Paneth cells and underlying mesenchymal cells to maintain stemness and proliferative capacity (Sato et al. 2009, Stzepourginski et al. 2017). A major advancement in organoid technology was the discovery that the mesenchymal niche was not required for stem cell maintenance as long as the appropriate growth factors were provided in enteroid media (Sato et al. 2009, Tekes et al.

2020). Many species have been successfully cultivated under these conditions, including human, mouse, dog, cow, sheep, horse, chicken, and pig enteroids (Powell & Behnke 2017, Tekes et al. 2020). The slow growth and early senescence of feline enteroids suggested to us and others (Powell & Behnke 2017, Tekes et al. 2020) that they might require growth factors not provided in traditional enteroid media. Tekes et al. used a human enteroid media formulation which delayed but did not rescue feline enteroid death. Powell & Behnke tested many additional growth factors for their ability to support cat enteroids, including FGF-2, -4, and -10, nicotinamide, IGF-1, PGE2, and Wnt-2b and Gremlin, but none rescued early growth arrest.

The same study saw that VIM+ mesenchyme-like cells were present in early cat enteroid passages. When these mesenchyme-like cells disappeared around passage 10, the cat enteroids began to decline. This suggested that mesenchymal cells provide necessary growth factors for long-term propagation of cat enteroids. We tested this idea by supplementing adult cat jejunal and ileal enteroids with mitomycin-C inactivated human foreskin fibroblasts (HFFs). We found that these HFF feeder cells were sufficient to support long-term growth of small intestinal jejunum enteroids and some ileum enteroids. HFF-supplemented small intestinal enteroids from cats and some mouse samples were also larger than non-supplemented enteroids, which may facilitate easier microinjection. RNA-seq data showed that enteroids and cell monolayers derived from enteroids are de-differentiated relative to intestine, which motivates future optimization of pro-differentiation conditions. Finally, transcripts of the canonical marker for Paneth cells were not detected in feline cell culture or intestine. Because these are the cells that normally support intestinal stem cells in other species, this raises an interesting possibility that the reason cat enteroids require HFFs is because they have no Paneth cells to provide survival-promoting growth factors.

## Results

### Selection of organoid media formulation

Our laboratory previously grew fetal and adult small intestinal organoids from the domestic cat (*Felis catus*) to study host-parasite interactions in *T. gondii* infection (Martorelli di Genova et al. 2019). However, the enteroids grew slowly and underwent growth arrest and cell death after 2-5 passages in culture. To determine whether we could optimize our media for cat enteroid growth, we consulted previous studies that used enteroids for infectious disease research (Table 1) (Powell & Behnke 2017, Co et al. 2019, Tekes et al. 2020). The human organoid media components nicotinamide, A-83-01, SB202190, and gastrin supported feline organoid growth for 10 additional passages in a previous study (Tekes et al. 2020). Powell & Behnke tested additional stem cell-promoting compounds, but they did not extend organoid survival. As such, we added A-83-01, SB202190, and gastrin to our original media formula (Martorelli di Genova et al. 2019).

**Table 1:**
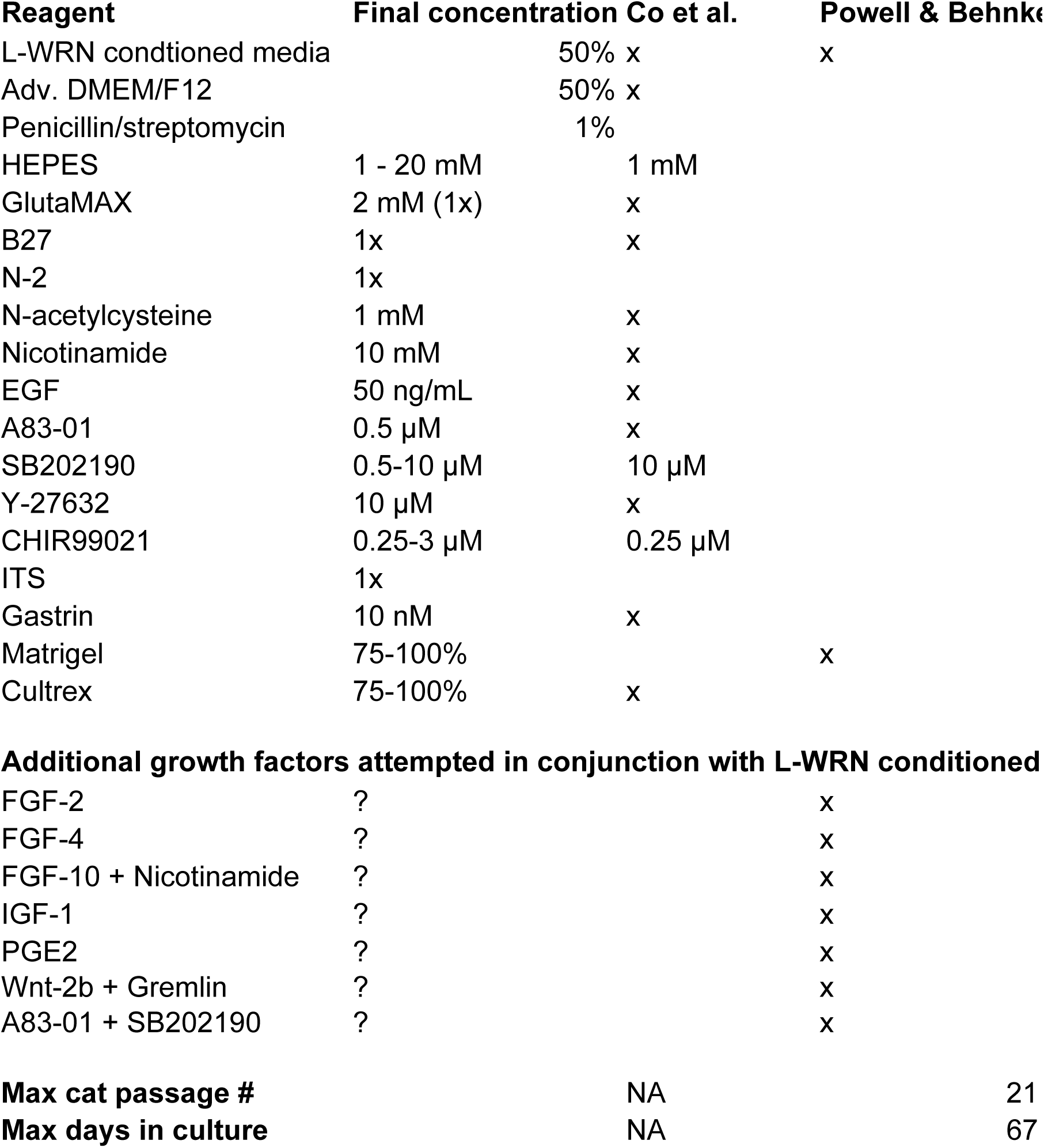
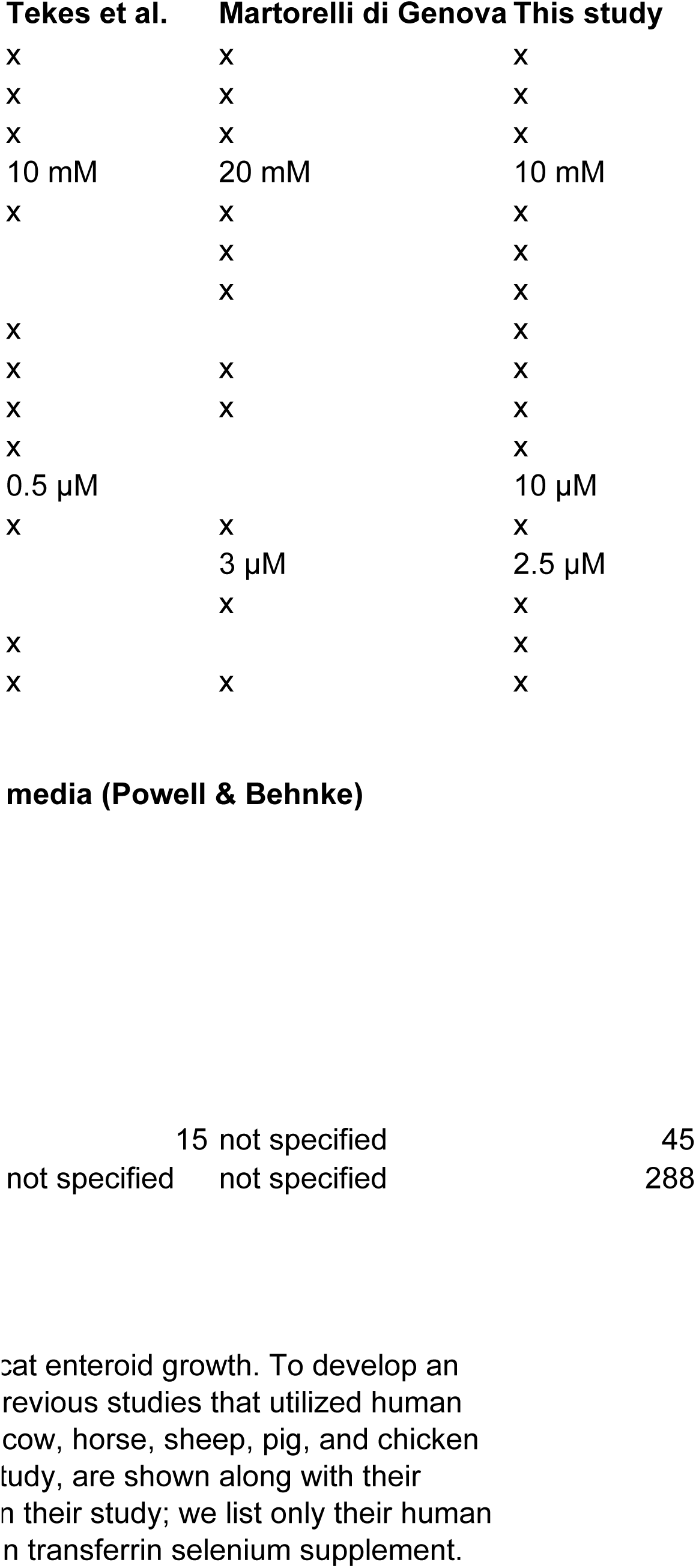
Organoid media formulations that enable cat enteroid growth. To develop an optimal growth media for feline enteroids, we consulted media formulations from previous studies that utilized human (Co et al. 2019), cat (Tekes et al. 2020), and a suite of species including cat, dog, cow, horse, sheep, pig, and chicken (Powell & Behnke 2017). Components used in each study, including the present study, are shown along with their maximum cat enteroid survival times. Tekes et al. list several media formulations in their study; we list only their human media formulation, which was most successful for cat enteroid culture. ITS = insulin transferrin selenium supplement.

Cats are unable to synthesize arachidonic acid from its precursor linoleic acid due to their lack of intestinal delta-6-desaturase (D6D) activity, which led us to test whether lipid supplementation would rescue feline enteroids from early cell death (Rivers et al. 1975, Sinclair et al. 1979). However, neither arachidonic acid, which promotes stem cell growth in murine organoids and gamma irradiated mice (Wang et al. 2020), nor a mixed lipid supplement (Sigma-Aldrich L0288) rescued the feline enteroids.

### Human foreskin fibroblasts enable long-term passage and increase the size of feline enteroids

Powell & Behnke (2017) noted that feline enteroids declined after the loss of VIM+ mesenchymal-like cells in their cultures. VIM is a marker for mesenchymal cells like fibroblasts, which support adult stem cells *in vivo* and in some models of intestinal cell culture such as induced pluripotent stem cells (iPSCs) and organoid-derived monolayers (Spence et al. 2011, Stzepourginski et al. 2017, Altay et al. 2019, Merenda et al. 2020). We tested whether human foreskin fibroblasts (HFFs) co-cultured with feline enteroids would enable their long-term survival. We chose HFFs because they are low maintenance, commercially available, and commonly used to culture *T. gondii*. To inactivate fibroblasts, we used mitomycin C (MMC), a double stranded DNA alkylating agent that is used to growth inactivate mouse embryonic fibroblasts for mouse embryonic stem cell culture (Bryja et al. 2006). After 2 hours of MMC treatment, HFFs were trypsinized and seeded into 24-well plates for enteroid seeding the next day (Fig. 1).

To pilot our fibroblast protocol, we isolated jejunal and ileal crypts from one adult feline (cat 0), cultured them in Matrigel and organoid media, and passaged them continuously with or without HFFs (Fig. 1). Enteroids grew out of all crypt samples at 10-20 enteroids per well at passage 0 (Fig. 2a). By passage 3, however, HFF-supplemented enteroids were more numerous than un-supplemented enteroids. By passage 4, just one un-supplemented well of jejunal and ileal enteroids remained, with 10 and 40 enteroids respectively. All remaining enteroids died by passage 5. In contrast, HFF-supplemented enteroids from both tissues increased to 40-250 enteroids per well by passage 4. Enteroids from both tissues maintained their high proliferative capacity out to passage 8.

**Figure 2:**
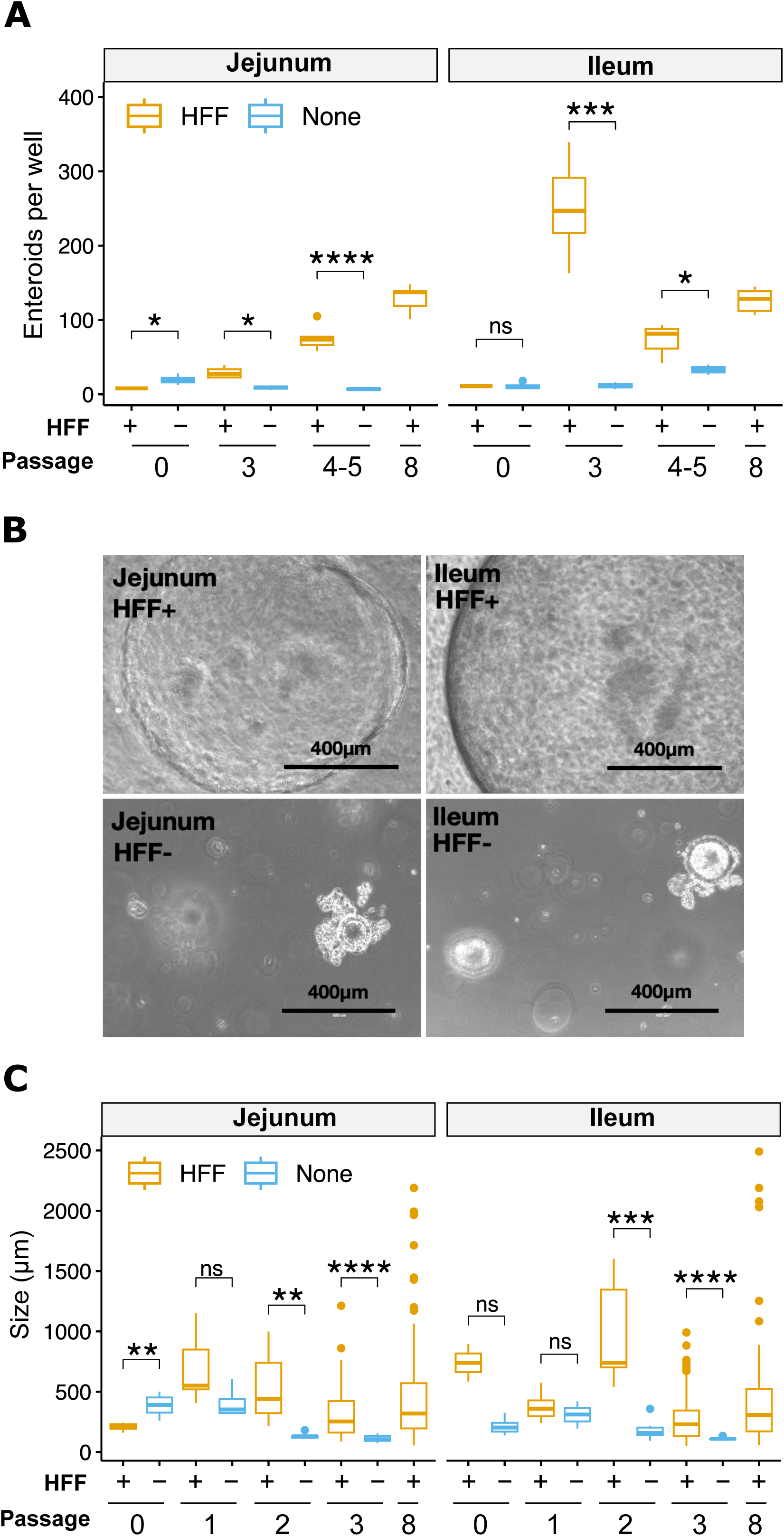
Fibroblasts promote early outgrowth of feline jejunum and ileum enteroids. Cat 0 enteroids were monitored from passage 0 to passage 8 to obtain enteroid counts per well (a), photos at 10x magnification using the EVOS imaging system (b), and enteroid diameters (c). All available wells were monitored, with at least 2 wells per condition until non-supplemented enteroids began to die at which point only one well remained. Enteroids were selected at random for photos and size measurements. *, p &lt; 0.05; **, p &lt; 0.01; ***, p &lt; 0.001; ****, p &lt; 0.0001, Student’s t-test.

We used the same enteroids from cat 0 to determine whether fibroblasts increase enteroid size (Fig. 2b-c). At passage 0, enteroids ranged in diameter from ∼200-700 µm with no obvious size difference between HFF supplemented and un-supplemented wells. Un- supplemented enteroids did not get any larger in subsequent passages, instead becoming smaller by passage 3 when they ceased proliferating. HFF-supplemented enteroids became larger over time, maintaining mean diameters of at least 200 µm from passage to passage with a subset of enteroids consistently reaching 1-2 mm diameters.

To test whether fibroblast feeder cells enhance enteroid survival in additional animals, we applied the HFF supplementation protocol to three other felines and tracked their survival (Table 2). All HFF-supplemented jejunal enteroids survived 126-288 days in culture (passage 23- 45), at which point they were preserved in liquid nitrogen. Ileal tissue was more variable. The ileal sample from cat 0 survived long-term while the samples from the remaining three felines died between passages 4 and 9. The ileal enteroids from cat 3 showed no signs of early senescence, but unfortunately they became contaminated with fungi at passage 9 and died shortly afterward. HFF-supplemented enteroids from cats 1-3 were more abundant and larger than un-supplemented enteroids (Fig. 3), including early ileum passages.

**Figure 3.**
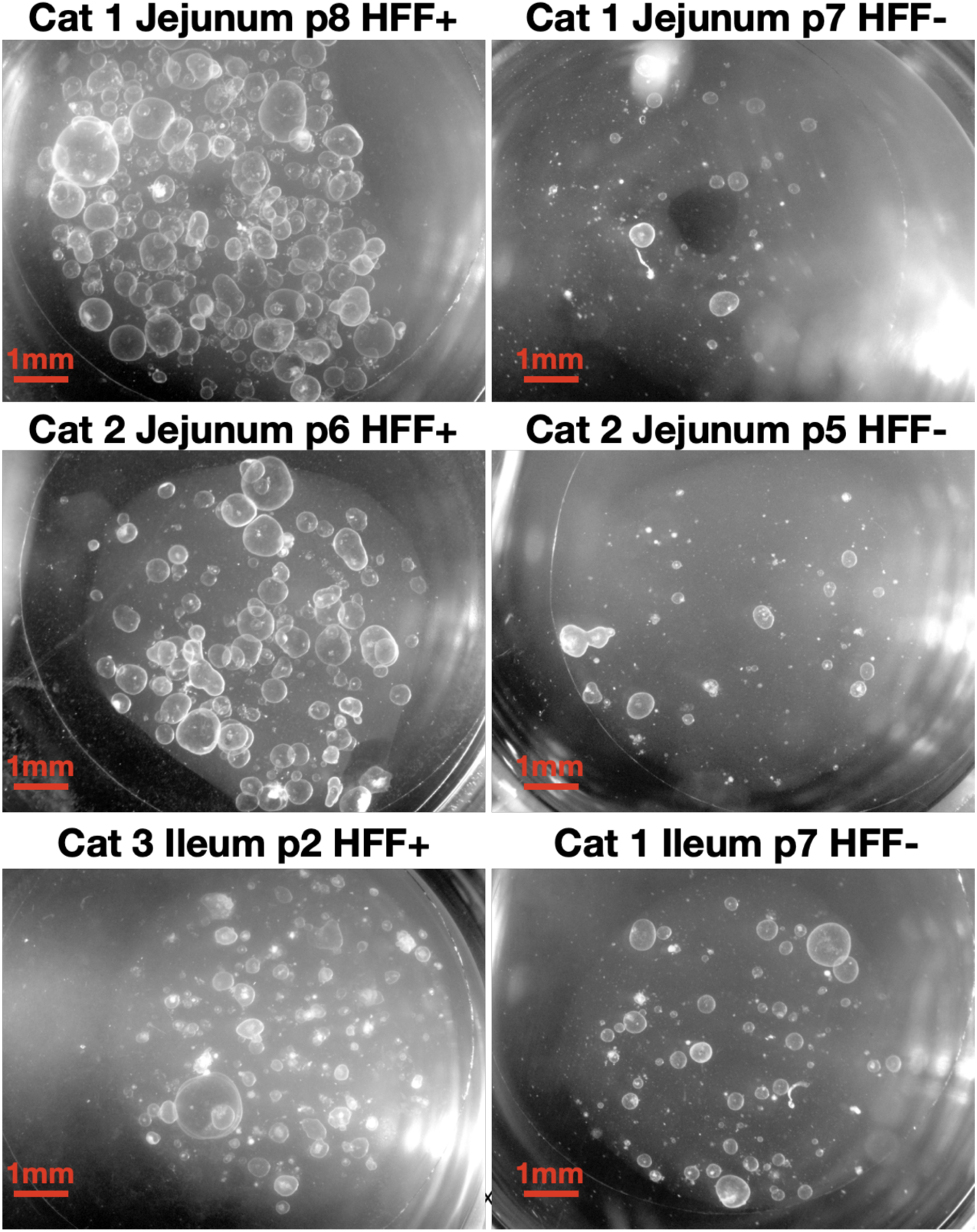
Fibroblasts increase the size and abundance of feline small intestinal enteroids. Representative photos of enteroids from cats 1, 2, and 3 grown with (left) and without (right) HFF supplementation. All photos were collected on the same date using Zeiss Zen software and a stereo microscope at 4x magnification.

**Table 2:** Cat enteroid survival times. Length of time in culture (passage number and total days) are listed for jejunum and ileum for each of the four cats used in this study. Samples used for fibroblast withdrawal experiments (cats 0 and 3) were offshoots of samples continuously passed with fibroblasts.

| Cat | Crypt Isolation Date | Tissue | Fibroblast Treatment | Passage Start | Passage End |
| --- | --- | --- | --- | --- | --- |
| 0 | 2/27/2023 | Jejunum | HFF | 0 | 45 |
| 0 | 2/27/2023 | Ileum | HFF | 0 | 45 |
| 0 | 2/27/2023 | Jejunum | None | 0 | 5 |
| 0 | 2/27/2023 | Ileum | None | 0 | 5 |
| 1 | 6/14/2023 | Jejunum | HFF | 0 | 29 |
| 1 | 6/14/2023 | Ileum | HFF | 0 | 7 |
| 1 | 6/14/2023 | Jejunum | None | 0 | 11 |
| 1 | 6/14/2023 | Ileum | None | 0 | 13 |
| 2 | 6/22/2023 | Jejunum | HFF | 0 | 26 |
| 2 | 6/22/2023 | Ileum | HFF | 0 | 4 |
| 2 | 6/22/2023 | Jejunum | None | 0 | 8 |
| 2 | 6/22/2023 | Ileum | None | 0 | 5 |
| 3 | 7/12/2023 | Jejunum | HFF | 0 | 23 |
| 3 | 7/12/2023 | Ileum | HFF | 0 | 9 |

Samples for Fibroblast withdrawal experiments
| Cat | Crypt Isolation Date | Tissue | Fibroblast Treatment | Passage Start | Passage End |
| --- | --- | --- | --- | --- | --- |
| 0 | 2/27/2023 | Ileum | None | 12 | 23 |
| 0 | 2/27/2023 | Jejunum | None | 17 | 44 |
| 0 | 2/27/2023 | Ileum | None | 17 | 44 |
| 3 | 7/12/2023 | Jejunum | None | 4 | 22 |
| 3 | 7/12/2023 | Ileum | None | 4 | 22 |

| Date Start | Date End | Days in Culture | Reason for Termination | Notes |
| --- | --- | --- | --- | --- |
| 2/27/2023 | 12/12/2023 | 288 | Froze |  |
| 2/27/2023 | 12/12/2023 | 288 | Froze |  |
| 2/27/2023 | 4/17/2023 | 49 | Died |  |
| 2/27/2023 | 4/17/2023 | 49 | Died |  |
| 6/14/2023 | 12/12/2023 | 181 | Froze |  |
| 6/22/2023 | 7/29/2023 | 37 | Died |  |
| 6/14/2023 | 8/24/2023 | 71 | Died |  |
| 6/14/2023 | 9/20/2023 | 98 | Died |  |
| 6/22/2023 | 12/12/2023 | 173 | Froze |  |
| 6/22/2023 | 8/3/2023 | 42 | Died |  |
| 6/22/2023 | 8/15/2023 | 54 | Died |  |
| 6/22/2023 | 7/26/2023 | 34 | Died |  |
| 7/12/2023 | 12/12/2023 | 153 | Froze |  |
| 7/12/2023 | 9/14/2023 | 64 | Died | died suddenly 9/14, culture had debris resembling fungal hyphae |
| 5/18/2023 | 8/15/2023 | 89 | Died |  |
| 6/27/2023 | 12/12/2023 | 168 | Froze |  |
| 6/27/2023 | 12/12/2023 | 168 | Froze | this sample was the only one with a budding phenotype |
| 8/8/2023 | 12/12/2023 | 126 | Froze |  |
| 8/8/2023 | 12/12/2023 | 126 | Died |  |

lture (passage number and amples used for fibroblast ith fibroblasts.

### Murine enteroids are less responsive than feline enteroids to fibroblast supplementation

Murine enteroids do not require fibroblast feeder cells for survival, but we reasoned that they might respond to growth-enhancing signals from fibroblasts. We tested whether HFF feeder cells would increase murine enteroid size or abundance (Fig. 4). HFF supplementation occasionally but not always increased enteroid counts in mouse jejunum and ileum samples (Fig. 4a). We did notice a significant increase in size for ileum enteroids (Fig. 4b), while jejunum enteroid size was less responsive to HFF supplementation. Overall, mouse and cat enteroids reached similar maximum sizes of 1-2 mm (Fig. 2c, Fig. 4a) and abundances of 200-300 enteroids per well (Fig. 2a, Fig. S4), with similar spherical (cystic) phenotypes (Fig. 3, Fig. 4b).

**Figure 4.**
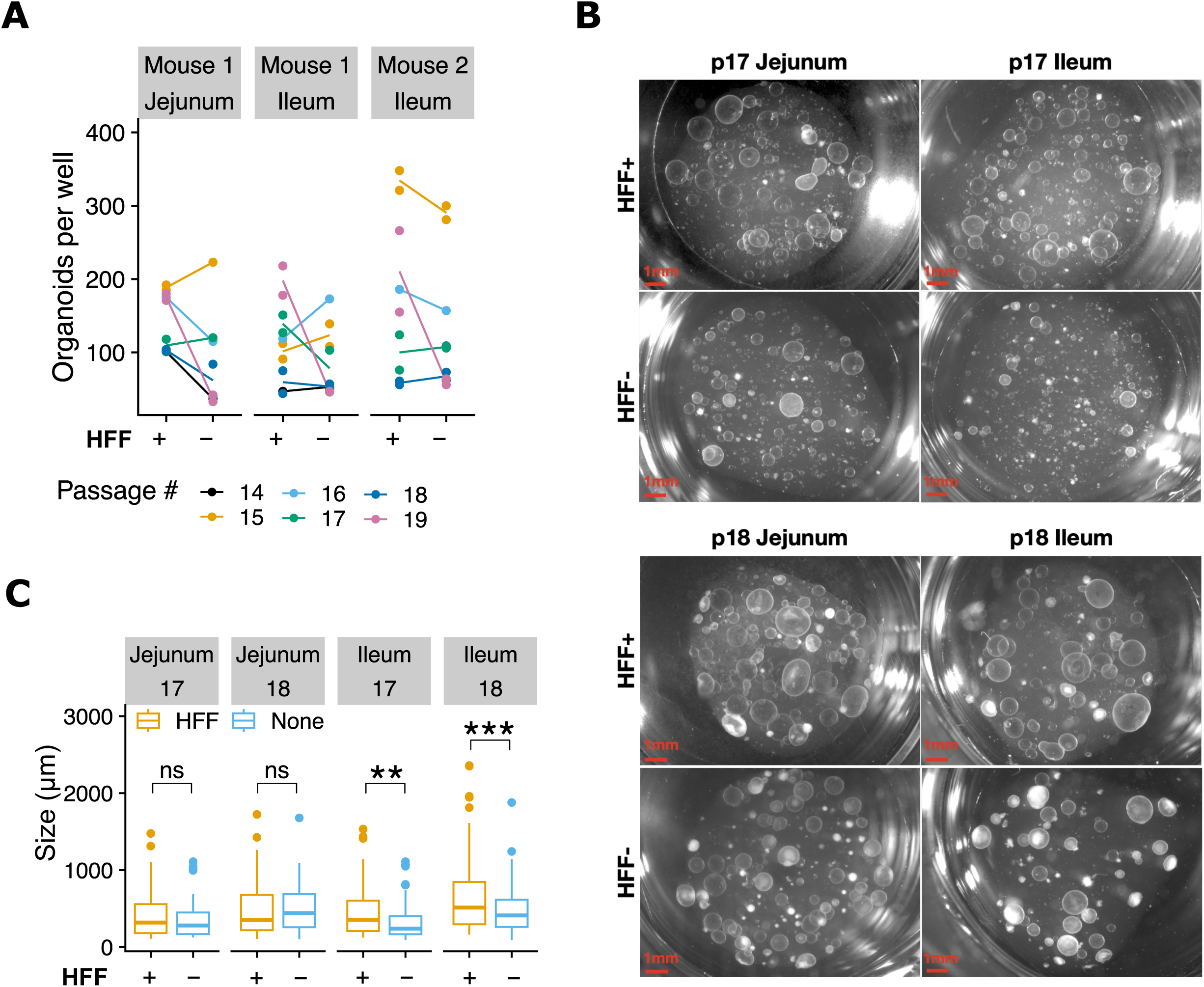
Murine enteroids are less responsive to fibroblasts than feline enteroids. (a) Per-well enumeration of murine jejunum (n=2 animals) and ileum (n=1 animal) enteroids throughout 6 passages of continuous maintenance with or without fibroblasts (n=1-2 wells per condition). No statistically significant differences were detected by paired t-test. **(b)** Representative phase images of enteroids from mouse 1 jejunum and ileum with and without HFF supplementation at passages 17 and 18. 4x magnification, scalebar = 1 mm. **(c)** The diameter of enteroids (n=25 per well, n=2 wells per condition for jejunum and n=4 wells per condition for ileum) were measured just before passaging for two consecutive passages, 17 and 18. *, p &lt; 0.05; **, p &lt; 0.01; ***, p &lt; 0.001; ****, p &lt; 0.0001, Student’s t-test.

### Long-term enteroid survival is possible following fibroblast withdrawal

Because fibroblast feeder cells require additional time and resources to maintain, we next tested whether we could stop fibroblast supplementation after enteroids were well-established (Table 2, Fig. 5a). We used samples from cat 0 and 3 for these experiments based on tissue availability, and compared un-supplemented samples to their parent samples that were continuously cultured with fibroblasts (Fig. 5b). For cat 3 (Table 2), we removed jejunum and ileum samples from fibroblasts at passage 4. Ileum enteroids remained abundant for around 15 passages but then dwindled until all were gone at day 126/passage 22. The jejunum enteroids appeared healthy through day 126/passage 22, at which point we preserved them in liquid nitrogen. An ileum sample from cat 0 was removed from fibroblasts at passage 12 and survived an additional 89 days/11 passages without fibroblast supplementation before enteroid death (Table 2). Enteroids were increasingly small and sparse leading up to their death (Fig. 5c). To test whether removing enteroids from fibroblasts at a later passage would enable longer survival, cat 0 jejunum and ileum were removed from fibroblasts at passage 17. Unlike the enteroids removed from fibroblasts at passage 12, both the jejunum and ileum enteroids survived at high abundance for an additional 168 days/27 passages prior to liquid nitrogen preservation (Table 2, Fig. 5d).

**Figure 5.**
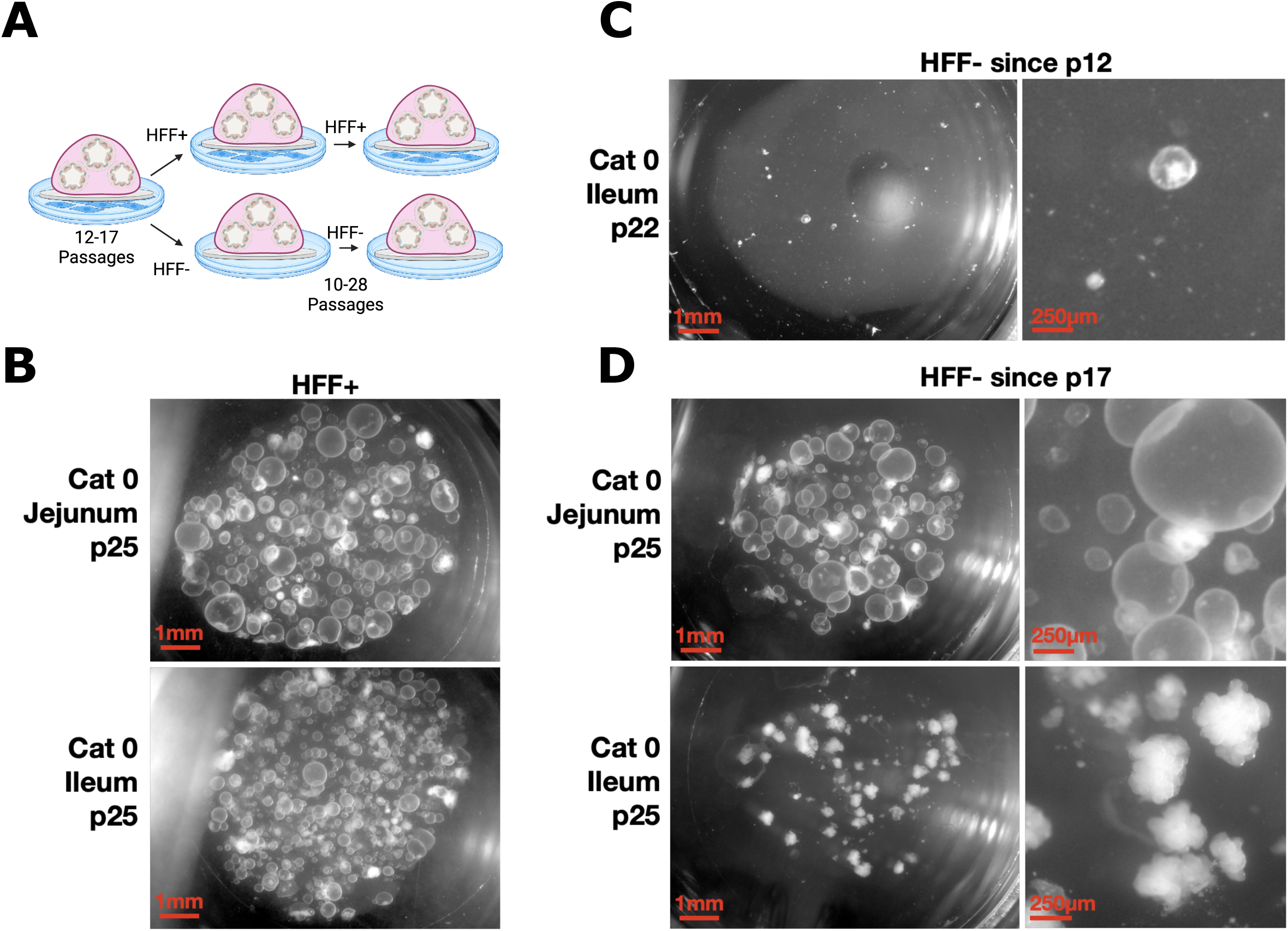
Long-term feline enteroid survival is possible following fibroblast withdrawal. (a) Schematic of experimental set-up. Cat 0 jejunum and ileum samples were initially grown on fibroblasts until fibroblast withdrawal at the indicated passage number, either 12 or 17. They were then split into two treatments, either continued fibroblast supplementation or continuous passage without fibroblast supplementation. Fig. 5a image created with Biorender.com. **(b)** Representative images of passage 25 cat 0 jejunum and ileum enteroids continuously passaged with HFFs. **(c)** Representative images of passage 22 cat 0 ileum enteroids that were grown without HFFs since passage 12. **(d)** Representative images of passage 25 cat jejunum and ileum enteroids that were grown without HFFs since passage 17. Note the budding phenotype of the ileum sample.

### Cat enteroids and their monolayers are de-differentiated with no Paneth cells

To begin to molecularly characterize the differentiation state of the feline enteroids, we extracted RNA from a small sample set and performed RNA-seq (Fig. 6). Our samples included one feline jejunum sample, one jejunal enteroid sample in Matrigel, and three jejunum organoid-derived monolayers (ODMs). We expected to see the highest levels of stem cell markers in the enteroid sample because Wnt3a is high in enteroid media, and because 3D suspension in Matrigel promotes a de-differentiated state (Co et al. 2019). Gut stem cell markers LGR5, SMOC2, and OLFM4 were indeed highest in the enteroid sample, and lower but present in intestine and ODMs. Enterocyte markers ALPI, FAPB2, and SI were highest in intestine. While ALPI was detectable in the enteroid and ODM samples, both FABP2 and SI reads were completely absent. Goblet cell marker MUC2 and enteroendocrine cell (EEC) marker CHGA were highest in intestine and lower in enteroids and ODMs. SYP, another EEC marker, was slightly elevated in jejunum relative to tissue culture samples. The Paneth cell marker LYZ was notably absent from all samples. In summary, enteroids and ODMs expressed higher levels of stem cell markers but had dramatically fewer differentiated cell types than *in vivo* jejunal epithelium.

**Figure 6.**
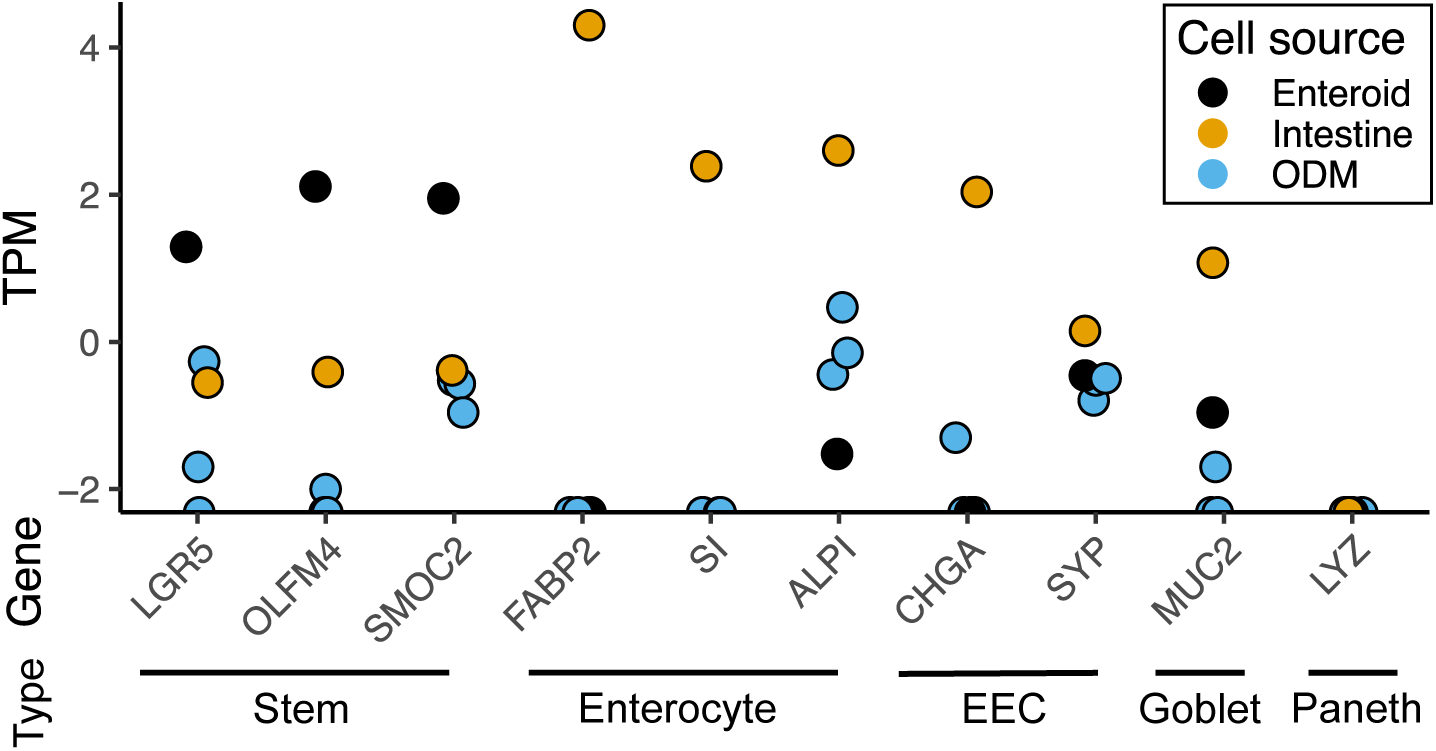
Molecular characterization of feline enteroids. mRNA transcripts were quantified by RNA sequencing of feline jejunum (n=1), feline jejunal enteroids (n=1), and monolayers derived from feline jejunal enteroids (n=3). Values for marker genes for stem cells (LGR5, OLFM4, SMOC2), enterocytes (FABP2, SI, ALPI), enteroendocrine cells (EEC) (CHGA, SYP), goblet cells (MUC2), and Paneth cells (LYZ) are reported as transcripts per million (TPM). ODM = organoid- derived monolayer.

## Discussion

We showed that feline small intestinal enteroids survive much longer than those in previous studies (Powell & Behnke 2017, Tekes et al. 2020) if they are supplemented with growth-inactivated human foreskin fibroblasts (HFFs). Jejunal enteroids were especially responsive to HFF supplementation with a survival rate of 100% (4/4 animals). Ileal enteroids were less responsive, with just 1/3 cultures surviving. Ileum enteroids tend be less proliferative than enteroids from higher in the gastrointestinal tract, which may warrant higher crypt seeding densities, more frequent changes of media and HFFs, or perhaps feline intestinal mesenchymal fibroblasts rather than HFFs. Mouse enteroids were less responsive to HFF feeder cells, though ileum enteroids were larger following HFF supplementation. This finding is consistent with the idea that mouse enteroids are already receiving the growth factors they need for optimal growth without HFF supplementation.

According to our RNA-seq data, feline jejunum enteroids and the monolayers derived from them were less differentiated than jejunum *in vivo*. Nearly all our feline enteroids were cystic in shape rather than budding, which is consistent with a de-differentiated state. Enteroid differentiation state is highly dependent on organoid media formulation. Our media, which most closely models human enteroid media with high levels of Wnt3a and stem cell-promoting SB202190, is known to maintain organoids in a de-differentiated, cystic state (Merenda et al. 2020). This de-differentiated state is problematic because a major benefit of enteroids is their ability to support both stem cells and terminally differentiated cell types in one biological unit. In fact, enteroid differentiation is necessary to support robust sexual development in the Apicomplexan parasite *Cryptosporidium parvum* (Heo et al. 2018, Wilke et al. 2019), making differentiation a high priority for our studies of *T. gondii* sexual development. Differentiation has been achieved for human colonoids by replacing Wnt3a and SB202190 with the growth factors IGF-1 and FGF-2. It may also be a useful starting point for feline enteroids. The organoid and parasitology fields have also established differentiation media formulations (Co et al. 2019, Holthaus et al. 2021) that may enhance feline enteroid differentiation. Finally, as shown in Figure 5, one of our cat 0 ileum samples developed budding structures consistent with differentiation after fibroblast withdrawal at passage 17. It maintained this morphology through passage 45 when we preserved the enteroids in liquid nitrogen, suggesting that fibroblast withdrawal alone may promote differentiation in some cases.

Paneth cells are important sources of stem cell-promoting factors in the small intestine of most mammals. Paneth cell ablation in mice results in enteroid death unless Paneth cell products like Wnt3a are provided in the media or by other epithelial or mesenchymal cells (Durand et al. 2012, Farin et al. 2012, van Es et al. 2019). We detected 0 transcripts of the Paneth cell marker LYZ in our feline intestine and cell culture samples (Fig. 6), which led us to wonder whether cat enteroids require fibroblasts because they lack Paneth cells. However, further investigation is necessary to determine whether Paneth cells are truly absent from cats. A previous study noted the absence of Paneth cells in cats, dogs, raccoons, and pigs (Gelberg 2014), but Paneth cells were later identified in pigs at low levels (Cui et al. 2023). While dogs lack expression of lysozyme, they do have Paneth-like cells that express other functional markers of Paneth cells (Chandra et al. 2019). Tekes et al. (2020) report low levels of LYZ gene expression in feline ileum tissue and enteroids, suggesting that Paneth cells may be present. In this study, we detected no lysozyme gene expression, but we only sequenced one jejunum sample due to RNA degradation in other intestinal samples. More studies are needed to resolve whether cats have Paneth cells, Paneth-like cells, or a Paneth cell-free epithelial compartment.

Feline enteroids are an important tool for studying viral and parasitic infections, but their unusually early growth arrest limits their usefulness. We showed that fibroblast feeder cells extend the lifetime of feline enteroids, especially in jejunum samples. In the future, we plan to optimize our protocols to robustly support ileum and colon enteriods. We will also test whether feline intestinal stem cells are dependent on fibroblasts to provide growth factors that are normally provided by Paneth cells in other species. Understanding how cats maintain their intestinal stem cell niche will be an important step toward accurate cell culture models of the feline intestinal environment for infectious disease research. We hope that replacing felines with feline enteroids will reduce the number of cats used in research. Our fibroblast supplementation method may also enhance the growth and survival of enteroids from other species whose intestinal tissue is slow-growing or prone to early senescence.

## Methods and Protocols

### Growth and Inactivation of Human Foreskin Fibroblasts

Human foreskin fibroblasts (HFF; ATCC, SCRC-1041) were cultured in high-glucose, no-glutamine DMEM supplemented with 10% fetal bovine serum, 2mM L-glutamine, 100 U/mL penicillin-streptomycin, and 10 mM HEPES at 37°C in a 5% CO-_2_ atmosphere. Confluent HFFs were then treated with mitomycin C (MMC) at 10 μg/mL and incubated at 37°C for 2-3 hours followed by 2 PBS washes to remove traces of MMC, dissociation with 20 μL/cm^2^ of 0.025% trypsin-EDTA for 3-5 minutes, then centrifuged for 3-5 minutes at 500 xg. The growth-inactivated HFFs were then seeded at 200,000 cells/ cm^2^ into 24-well plates and used as feeder cells for up to 8 days.

### Animals

Mice were treated in compliance with the guidelines set by the Institutional Animal Care and Use Committee (IACUC) of the University of Wisconsin School of Medicine and Public Health (protocol #M005217). Cats were euthanized for reasons unrelated to this specific research and were treated in compliance with the guidelines set by IACUC protocols.

### Organoid Media

Conditioned media containing Wnt3a, R-Spondin and Noggin was produced from the L-WRN cell line (ATCC CRL-3276) cultured in Advanced DMEM/F12, 20% fetal bovine serum, 2mM Glutamax, 10 mM HEPES, and 100 U/mL penicillin/streptomycin, as previously described (Miyoshi & Stappenbeck 2013). L-WRN conditioned media was frozen at -20°C in 25 mL aliquots until time of use. Media for culturing enteroids contained 40% Advanced DMEM/F12, 50% L-WRN conditioned media, 2 mM GlutaMAX, 10 mM HEPES, 1X B27, 1X N2, 1X insulin/transferrin/selenium, 50 ng/ml human EGF, 10 mM nicotinamide, 2.5 μM CHIR-99021, 10 μM Y-27632, 0.5 μM A-83-01, 10 μM SB202190, 10 nM human gastrin (all components listed in Table 1) and sterile filtered. Media was kept at 4°C in the dark and used for up to 2 weeks before discarding.

### Crypt Isolation

3 cm jejunum and ileum sections were obtained from 4 young cats and 2 mice. The section’s lumens were washed with a cold PBS solution (phosphate buffered saline (PBS), 1X penicillin/streptomycin, 25 μg/mL gentamicin), cut open and cut into 1 cm pieces. The small sections were then washed 3 times by gentle rotation for 30 seconds in cold PBS solution. As a final wash, sections were rotated in PBS solution for 20 minutes at 4°C. Sections were then placed in 3mM EDTA in PBS on wet ice for 30 minutes with no agitation followed by another 30 minute incubation in ice-cold 3mM EDTA in PBS with tube rotation at 4°C. Sections were shaken vigorously for 30 seconds to release the crypts, inspected by light microscope, and passed a 70 μm cell strainer to remove villi. Crypts were spun at 250xg for 10 minutes at 4°C, supernatant removed, and resuspended in 250 – 500 µL enteroid media. We recommend diluting approximately 10 µL enteroid solution to 40 µL Matrigel for plating. Dilution may need to be determined empirically; the goal is to maximize enteroid outgrowth. For plating, begin at Step 10 of the *‘Culturing Enteroid Plugs’* protocol below.

### Culturing Enteroid Plugs

1. Enteroids should be passaged every 3-7 days or when enteroid cores become dense with necrotic cells. MMC-treated HFFs are stable for 6-8 days in enteroid media, though they decline in health either due to MMC treatment or to enteroid media. Note: If enteroids grow slowly, they can be carefully and gently moved to a new HFF monolayer by grasping their coverslip with forceps. We recommend practice with this technique before working with valuable enteroids.
2. Visually assess enteroids to determine passage density. Either count enteroids and plan to seed 50-100 enteroids per new well for 24-well plates or passage at 1:3-1:4 as a starting point. Adjust ratios for enteroid density and expect small, budding, or dying enteroids to be less proliferative than large spherical enteroids without necrotic cores.

E.g. for the enteroids depicted in Figure 3 (starting at the top left with the Cat 1 Jejunum p8 HFF+ sample and moving counter-clockwise), we would recommend passaging enteroids at ratios of 1:8, 1:6, 1:4, 1:2, 1:1, and ending with 1:1 for the Cat 1 Jejunum p7 HFF- sample. We would passage the Cat 0 p25 Ileum HFF- sample in Figure 5c at 1:2 or 1:3.

1. 3. Preparation: Unsure HFFs in HFF plates are confluent. UV treat non-fibroblast plates for 15 minutes and pre-warm them at 37°C. Pre-chill a tabletop microcentrifuge to 4°C. Pre-chilled PBS with 1% FBS should be kept on wet ice. Pre-warm enteroid media to 37°C. Thaw Matrigel aliquots on wet ice. Note on glass coverslips: HFF-supplemented enteroids need to be plated on glass coverslips to prevent their direct contact with the HFF monolayer. If direct comparisons between HFF-supplemented and non-supplemented enteroids are required, we recommend adding and UV-ing glass coverslips to non-fibroblast plates for consistency. We autoclave glass coverslips and store them in a glass Petri dish with taped lid. Other notes: In our hands, adding 1% FBS to PBS aids in separating enteroids from Matrigel, especially cat enteroids. Our protocol assumes use of 24-well plates; adjust volumes of Matrigel, enteroids, trypsin, and PBS if using other plates.
2. 4. Beginning passage: Remove media from enteroids and place the 24-well plates on wet ice to begin dissolving Matrigel. Add 250-500 µL ice-cold PBS to each well. Dislodge plugs with pipette tip, avoiding MMC-treated fibroblast monolayer if applicable. Aspirate and dispense the PBS-containing dislodged plugs twice to aid in dissolving Matrigel.
3. 5. Break open enteroids and further disrupt Matrigel by slowly aspirating liquid through a 27G needle attached to a 1-, 3-, or 5-mL syringe, avoiding bubbles. Dispense back into well. Repeat the aspiration step, then dispense into a 1.7 mL Eppendorf tube. Pool up to 4 wells of enteroids per tube to aid in pelleting. Centrifuge at 2500 xg for 10-15 seconds at 4°C. Aspirate PBS and as much Matrigel as possible without disturbing the underlying cell pellet. Repeat PBS, spin, and supernatant removal once more.
4. 6. Trypsin treatment was used as necessary for mouse enteroids when enteroids were too small to lyse with a syringe. Trypsin treatments were done with 250-300 µL of 0.05%- 0.25% Trypsin for 5 minutes at 37°C with a shake step halfway through incubation. Trypsin was inactivated with enteroid media followed by 5-10 syringes with a 25G or 27G needle. Enteroids were pelleted at either 500xg for 5 minutes or 2500xg for 10-15 seconds at 4°C and media aspirated. Note: In our hands, the 1% FBS in PBS is compatible with trypsin at the trypsin concentrations listed.
5. 7. Resuspend enteroid pellets from step 3 or step 4 in enteroid media and spin again at 2500xg for 10-15 seconds at 4°C to remove residual trypsin or PBS. Remove supernatant.
6. 8. Resuspend enteroid pellets in enteroid media at 10 µL per enteroid plug. E.g. if you pooled 2 high-density plugs to be split at 1:8, dilute in 160 µL enteroid media. Note: we recommend enteroid dilution at this step, not earlier. Enteroids pellet best when they are abundant.
7. 9. Prepare an Eppendorf tube for each tissue with 10 µL enteroid solution and 40 µL Matrigel for each new well. Mix well, pipetting slowly to avoid bubbles.
8. 10. Remove pre-warmed plates from 37°C just prior to plating. Aspirate media from HFF- containing wells and place glass coverslips directly onto HFF monolayers using sterile forceps or Pasteur pipette attached to vacuum apparatus. Note: to avoid shearing the monolayer, take care not to move the glass coverslip once it is placed onto the cells. We use glass coverslips because Matrigel in direct contact with the moist HFF monolayer spreads out rather than retaining its dome shape. Coverslips also aid in preventing HFFs from transferring from one passage to the next.
9. 11. Add 50μL drops of Matrigel/enteroid to the glass coverslip in each well, avoiding contact with the HFF monolayer. Note: Matrigel is viscous and must be pipetted ice cold with a pre-chilled pipette tip to evenly distribute enteroids within and across wells. Pipette slowly and do not use the second stop to avoid bubbles.
10. 12. Quickly move enteroids to 37°C incubator to set Matrigel until it is does not jiggle when plate is tapped (5-10 minutes). Note: In our hands, if Matrigel was not set quickly, cat enteroids tended to sink and attach to glass or plastic, where the spheroids opened into monolayers and died. Avoid tipping or bumping the plate.

Add 500 µL enteroid media and return to incubator. Change enteroid media every 2-3 days.

### Enteroid enumeration and size measurements

Low-passage (passages 0-8) enteroids from cat 0 were imaged and measured at 10x magnification using an EVOS FL Auto imaging system. Photos of higher-passage cat enteroids and mouse enteroids were collected using a Zeiss Stemi 2000-C stereo microscope at 4x magnification for counting and measuring (Zeiss Zen software). Some enteroids were too small to reliably count and Matrigel plugs that leaked beyond the edges of coverslips during plating were not counted.

### Preparing and Culturing Enteroid Monolayers

1. Prepare a working solution of 1.0 mg/mL rat tail collagen on ice using the following volumes per 1 mL of solution: 100 µL 10X PBS, 4.6 µL 0.5 M NaOH, 795 µL sterile water, and 101 µL of 9.9 mg/mL rat tail collagen. Recommended range of working stock collagen concentrations: 0.5 - 2.0 mg/mL. Collagen stock concentrations vary by lot. Volume of NaOH should be adjusted to the volume of collagen added, at a ratio of 1:0.023 collagen:NaOH. Reagents should be added in the order listed.
2. Add enough collagen solution to wells to completely coat bottoms of wells. After even coating, remove collagen with a pipette and transfer to the next coverslip-containing well until all coverslips are coated. Cure collagen in a 37°C humidified incubator with CO_2_ for 3-6 hour(s).
3. UV treat collagen-coated wells with UV for 15 minutes.
4. Cover collagen with cell culture media (usually HFF culturing media) then incubate overnight in a 37°C humidified incubator with CO_2_.
5. *Add enteroids*: Follow the *Culturing Enteroid Plugs* protocol above through the trypsinization step. Wash once with enteroid media and centrifuge 2500xg for 10-15 seconds to remove residual trypsin. Resuspend cell pellets in enteroid media and distribute evenly among wells. Add more media immediately after cell plating for final volumes ∼500 µL for 24-well plates or ∼1.5 mL for 12-well plates. Recommended dilution: 3-4 (24-well plate) wells of monolayers per 1 (24-well plate) well enteroids.
6. Change enteroid media every 2-4 days. Note: Enteroid monolayer attachment and health is more sensitive to cold and desiccation than organoids in Matrigel. Use fully pre-warmed media for media changes, perform media changes quickly, and consider placing a warm plate between the enteroid plate and laboratory surfaces.
7. Monolayers can be plated on collagen-coated glass coverslips for imaging. Simply add coverslips prior to adding collagen, ensure the coverslip is completely coated, and press the coverslip to the bottom of the plate using a sterile pipette tip. If cell attachment is poor, consider other attachment matrices or alternative coverslips (e.g. gelatin, entactin-collagen-laminin, poly-L-lysine coated coverslips).

### Generation of RNA and RNA sequencing

Cell pellets were digested in Trizol and stored at -80°C until time of extraction. 50-150 mg of tissue samples were flash frozen in solvent-resistant screw-cap tubes containing 0.1 mm zirconia/silica beads (BioSpec Products 11079101z) and 1 large stainless steel bead (BioSpec Products 11079132ss), thawed in Trizol, and lysed for 3 minutes on a bead beater on high speed at room temperature. Phenol-chloroform extraction was performed, and samples were treated with DNAse I prior to TapeStation quality control and library preparation. We used the University of Wisconsin—Madison Biotechnology Center’s Gene Expression Center Core Facility (research resource identifier [RRID]: SCR_017757) for RNA quality control and library preparation, and the DNA Sequencing Facility (RRID: SCR_017759) for sequencing and demultiplexing of reads. Libraries were prepared with poly(A) enrichment (Illumina TruSeq stranded mRNA). Samples were sequenced on 2 lanes of a 2 × 150-bp Flowcell (NovaSeq S4). On average, a total of 83.24 million reads (minimum, 63.2 million; maximum, 108.8 million) paired-end 150-bp reads per sample were generated. The quality of the reads was determined using FastQC (Andrew & Bittencourt 2010), a threshold of 34 was selected, and only reads that met the threshold were used for further analysis. We trimmed the data to remove low-quality reads using Trimmomatic (Bolger et al. 2014) Reads were mapped to the cat genome *Felis catus* (NCBI RefSeq Assembly F.catus_Fca126_mat1.0), was conducted using STAR (Spliced Transcripts Alignment to a Reference program) (Dobin et al. 2013, Dobin & Gingeras 2015) using default parameters. Quantification of mapped reads and the generation of a counts table were conducted using RSEM (Li & Dewey 2011). Counts were imported into RStudio for data analysis and figure preparation using R.

### Statistical Analysis

Figure preparation and statistical analyses were performed in RStudio using the R programming language.

## Acknowledgments

The authors thank members of the Knoll laboratory for helpful feedback and discussions. This work was supported by a National Institutes of Health National Institute of Allergy and Infectious Diseases R01AI144016-01 (L.J.K.), and a Ruth L. Kirschstein Postdoctoral Individual National Research Service Award from the National Institutes of Health National Institute of Allergy and Infectious Diseases F32 AI172084 (N.D.H.). The content is solely the responsibility of the authors and does not necessarily represent the official views of the National Institutes of Health.

